# No Evidence for Systematic White Matter Correlates of Dyslexia and Dyscalculia

**DOI:** 10.1101/259788

**Authors:** David Moreau, Anna J. Wilson, Nicole S. McKay, Kasey Nihill, Karen E. Waldie

## Abstract

Learning disabilities such as dyslexia, dyscalculia and their comorbid manifestation are prevalent, affecting as much as fifteen percent of the population. Structural neuroimaging studies have indicated that these disorders can be related to differences in white matter integrity, although findings remain disparate. In this study, we used a unique design composed of individuals with dyslexia, dyscalculia, both disorders and controls, to systematically explore differences in fractional anisotropy across groups using diffusion tensor imaging. Specifically, we focused on the corona radiata and the arcuate fasciculus, two tracts associated with reading and mathematics in a number of previous studies. Using Bayesian hypothesis testing, we show that the present data favor the null model of no differences between groups for these particular tracts—a finding that seems to go against the current view but might be representative of the disparities within this field of research. Together, these findings suggest that structural differences associated with dyslexia and dyscalculia might not be as reliable as previously thought, with potential ramifications in terms of remediation.

## 1. Introduction

Reading and mathematics are essential skills mastered by most children at a young age, yet despite adequate intelligence and educational opportunities, many individuals exhibit difficulties that often continue well into adulthood. Dyslexia is a common neurodevelopmental disorder characterized by severe difficulties in reading and spelling that are typically difficult to address via usual remedial efforts (Eden & Zeffiro, 1998; S. E. Shaywitz et al., 1998; S. E. Shaywitz & Shaywitz, 2005; Vellutino, Fletcher, Snowling, & Scanlon, 2004). Dyscalculia, often thought of as the mathematical counterpart to dyslexia, is characterized by a severe and persistent impairment in mathematics, including difficulties using number and quantity, simple arithmetic and counting (Kucian et al., 2014; Reigosa-Crespo et al., 2011; Wilson et al., 2015). Although approximately one student in every class will exhibit mathematics-related difficulties, dyscalculia remains relatively under-researched in comparison to dyslexia (Badian, 1999; Barbaresi, Katusic, Colligan, Weaver, & Jacobsen, 2005; Gersten, Clark, & Mazzocco, 2007; Kucian et al., 2014). For reasons that remain to be fully understood (Rubinsten, 2009), both disorders often co-occur—given the presence of one, the prevalence of the other is approximately 40% (Wilson et al., 2015). A large body of research has examined whether these learning disorders have underlying anatomical, as well as a functional, bases. Functional and structural imaging techniques have been used to compare the brains of typical achievers with those of individuals who exhibit either math difficulties, reading difficulties, or both—documenting a range of differences in functional activation and white matter connectivity.

At the functional level, reading is a complex multi-faceted task that relies on a widespread left lateralized network. This network includes several temporo-parietal regions and temporo-occipital regions, together with regions within the inferior frontal gyrus, including Broca’s area (Fiez & Petersen, 1998; Pugh et al., 2001). Specifically, co-activation of subregions within these areas has been proposed to underlie the direct process of converting word-form perception to word meaning, otherwise known as the lexicosemantic route, whereas distinct subregions of the same areas are thought to be involved in the graphophonological, or indirect, route (Jobard, Crivello, & Tzourio-Mazoyer, 2003). In dyslexic individuals, this reading network is typically impaired (Brunswick, Mccrory, Price, Frith, & Frith, 1999; Shaywitz et al., 2002; Shaywitz et al., 1998), such that activation in both the temporo-parietal and temporo-occipital areas is significantly reduced, while right-hemisphere activation is increased (Brunswick et al., 1999; B. A. Shaywitz et al., 2002; S. E. Shaywitz et al., 1998). This increased activation in the right hemisphere is often thought to reflect a compensation effect resulting from the left-hemisphere hypoactivation (Shaywitz et al., 2002). Consistent with this idea, a connectivity study in a population of children with reading difficulties reported modulation of both the left inferior frontal gyrus and the left inferior parietal lobule, compared to typical readers (Cao, Bitan, & Booth, 2008).

Carrying out mathematical tasks engages a neural network linking the frontal and parietal regions, which partially overlaps with the reading network (Ansari et al., 2008; Dehaene, 1999). The importance of this network in mathematical ability was highlighted in a study by Emerson and Cantlon (2012), who found that fronto-parietal connectivity in 4-11 year old children was correlated with scores on a mathematical ability test, independent of IQ. In addition, individuals with dyscalculia typically show reduced neural activity in both parietal and frontal areas when performing mathematical tasks (Kucian et al., 2006), although the specific subregions vary considerably between studies. A few connectivity studies have suggested that connectivity of the fronto-parietal network is altered in dyscalculics, yet the nature of the abnormality remains unclear (Butterworth et al., 2005). Two functional connectivity studies have shown hyperconnectivity between the intraparietal sulcus (IPS) and frontal regions in children with dyscalculia compared to typically developing children, whereas Kucian and colleagues (2014) showed reduced structural connectivity in the superior longitudinal fasciculus (SLF).

The structural correlates of underlying differences in functional connectivity have been examined using Diffusion Tensor Imaging (DTI), which measures the water diffusion within brain tissue, taking advantage of the fact that the structural properties of white matter result in anisotropic (or directional) diffusion of water molecules (Beaulieu, 2002; Beaulieu, 2009; Soares, Marques, Alves, & Sousa, 2013). The directionality of water diffusion, thought to be a good estimate of white matter integrity (Beaulieu, 2002; Beaulieu, 2009), can be probed via fractional anisotropy (FA)—high FA typically occurs where myelinated axons are dense and have directional coherence, as underlying structure constrains the direction of movement.

In dyslexic readers, FA has been found to be reduced in the temporo-parietal region in comparison to typical readers (Deutsch et al., 2005; Klingberg et al., 2000; Vandermosten, Boets, Wouters, & Ghesquière, 2012). In addition, several studies have identified regions showing an association between reading ability and FA, including positive correlations in the temporo-parietal and frontal areas and negative correlations in the posterior corpus callosum (Niogi & McCandliss, 2006; Odegard, Farris, Ring, McColl, & Black, 2009; Rimrodt, Peterson, Denckla, Kaufmann, & Cutting, 2010). A number of white matter tracts have been linked with reading, including the corona radiata (CR), arcuate fasciculus (AF)— a subregion of the SLF (Vandermosten, Boets, Wouters, et al., 2012)—and the corpus callosum (Catani & Thiebaut de Schotten, 2008; Niogi & McCandliss, 2006; Odegard et al., 2009). A meta-analysis of nine DTI studies in both children and adults conducted by Vandermosten et al (2012) suggested that two white matter tracts, the AF and the CR, show reduced FA in dyslexia. Taken together, these findings suggest that dyslexic readers may exhibit reduced connectivity between areas that typically correlate with greater reading ability.

A few studies have investigated white matter tracts associated with mathematics; the most salient reported results are correlations between FA performance on a range of mathematical tasks in the AF, superior CR and, to a lesser extent, in the inferior longitudinal fasciculus (ILF; Beek, Ghesquière, Lagae, & De Smedt, 2014; Klein, Moeller, Glauche, Weiller, & Willmes, 2013; Matejko, Price, Mazzocco, & Ansari, 2013; Rykhlevskaia, Uddin, Kondos, & Menon, 2009; van Eimeren, Niogi, McCandliss, Holloway, & Ansari, 2008). Less research has focused on the white matter integrity correlates of dyscalculia. Kucian et al. (2014), using a whole-brain analysis, found that children with dyscalculia show reduced FA in the posterior SLF, whereas in a probabilistic tractography study, Rykhlevskaia et al. (2009) found that the tracts most implicated were the ILF and the inferior fronto-occipital fasciculus.

Despite the high rate of comorbidity between dyslexia and dyscalculia, there is no published research that has investigated whether the white matter integrity differences found in one of the two disorders vary as a function of the other disorder also being present. A recent study used DTI to examine mathematical difficulties in dyslexic children (Koerte et al., 2016). Individuals with dyslexia are known to experience particular mathematical difficulties with verbal arithmetic, likely related to impaired phonological retrieval processes (Boets & De Smedt, 2010). In this study, arithmetic efficiency showed a greater correlation with FA in children with dyslexia vs. controls in a large number of white matter tracts, including the bilateral ILF and SLF (Koerte et al., 2016).

The present study aimed to investigate the relationship between white matter connectivity and comorbidity of dyscalculia and dyslexia, via Tract-Based Spatial Statistics (TBSS) and probabilistic tractography. TBSS is a whole brain data-driven analysis that was used in the present study to investigate whether between-group differences in FA could be observed in *any* white matter regions. Probabilistic tractography allowed us to investigate FA differences within *specific* tracts hypothesized from previous literature to be important for both reading and mathematics. Together these two techniques enabled complementary whole-brain and ROI approaches to white matter connectivity. The seed regions selected for tractography were located within the CR and AF, based on previous literature linking these tracts to reading and mathematical difficulties (Vandermosten et al., 2012; van Eimeren et al., 2008; Matejko et al., 2013).

## 2. Materials and Methods

### 2.1 Participants

Subjects were recruited as part of the Auckland Comorbidity Study (Waldie, Wilson, Roberts, & Moreau, 2017; Wilson et al., 2015). From 85 adults participating in the larger behavioral study, 47 were selected for the current study. Two subjects were removed due to problems with data collection. The final sample (n = 45) included 11 dyslexics, 11 dyscalculics, 12 comorbids and 11 controls, all right-handed. Initial screening included the administration of the Wechsler Abbreviated Scale of Intelligence (WASI), standardized tests of reading, spelling and mathematics (Woodcock Johnson Word ID and Word Attack, and the Wide Range Achievement Test spelling and mathematics), and a clinical assessment for ADHD using the Adult Self Report Scale (ASRS) as a screener, followed by a diagnostic clinical interview for participants with high ASRS scores. Participants with dyslexia and dyscalculia participants reported a history of reading/mathematic difficulties, together with current difficulties. The criteria for dyslexia were at least one score ≤ 25th percentile on one of the reading and spelling tests (WJ Word ID and Word Attack, and WRAT spelling), plus at least one other of these scores ≤ 50th percentile. The criterion for dyscalculia was a score ≤ 25th percentile on the WRAT mathematics. Exclusion criteria for all groups included the presence of a neurological disorder (except mild depression or anxiety), a history of major head injury or non-standard schooling, English as a second language, vision or hearing impairment, Full Scale Intelligence Quotient (FSIQ) < 85, clinical diagnosis of ADHD, and fMRI contraindications. Further details regarding the screening procedure are available in Wilson et al. (2015). All participants gave informed consent with approval from The University of Auckland Human Participants Ethics Committee. Sample characteristics are shown in Table 1.

**Table 1.**
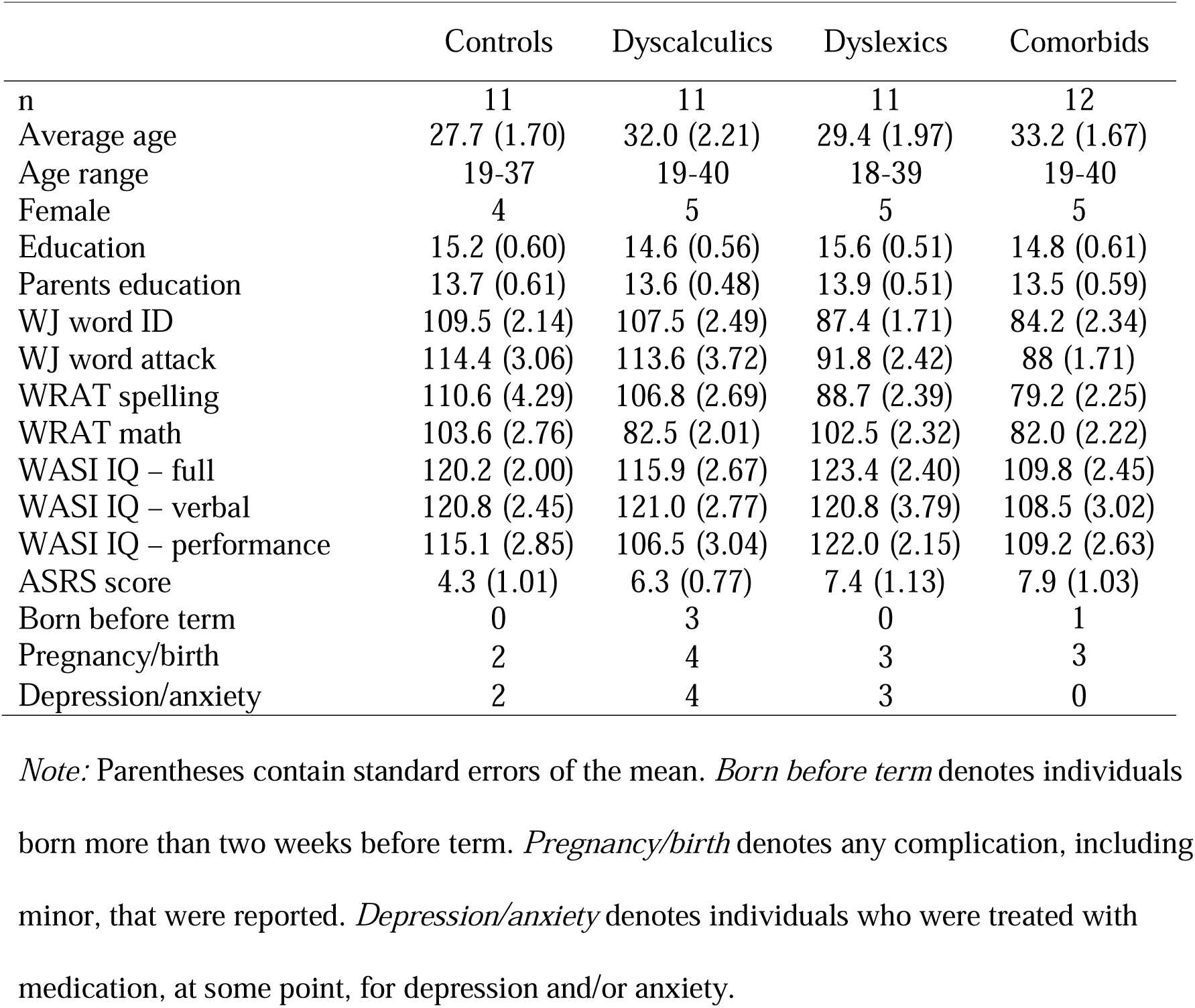
Sample characteristics, broken down by groups (controls, dyscalculics, dyslexics, and comorbids).

### 2.2 Data acquisition

Diffusion data was acquired on a 1.5-T Siemens Avanto scanner (Erlangen, Germany) using a single shot spin-echo EPI sequence (TR 6601; TE 101; FOV 230; voxel size 1.8×1.8×3.0mm) in 30 non-collinear directions. Images were obtained with a diffusion weighting of b = 1000s/mm^2^ and a reference image with a diffusion weighting of b = 0s/mm^2^. The sequence was applied twice and averaged to increase the signal to noise ratio. T-1 structural volumes were acquired using a fast low angle shot (FLASH) sequence (TR 11; TE 4.94; FOV 256; 176 sagittal slices; voxel size 1x1x1mm).

### 2.3 Data preprocessing

Images were processed using FMRIB’s software library (FSL) version 5.0 (Oxford Centre for Functional MRI of the Brain [FMRIB], UK). Images were converted from DICOM files to the appropriate NIFTI format using MRICRON (https://www.nitrc.org/projects/mricron). FSL’s brain extraction tool (BET) was used to remove any non-brain tissue and create a brain mask (Smith, 2002). Eddy current distortions and motion artifacts were corrected for using the eddy current correction tool (Jenkinson & Smith, 2001). The no-diffusion image of each subject was then used to create a binary brain mask. Lastly, for each subject, we generated FA maps by fitting the diffusion tensor model using the DTIFIT tool provided by FSL (Behrens et al., 2003; Behrens, Berg, Jbabdi, Rushworth, & Woolrich, 2007). All subjects were visually inspected to ensure preprocessing was satisfactory. Preprocessing for five subjects (11% of the sample) had to be manually corrected given that the default parameters resulted in incorrect segmentation of the brain from the skull.

### 2.4 TBSS

We used a whole-brain approach to look at general white matter differences in preprocessed FA values. Tract-based spatial statistics (TBSS) allows the implementation of a unique nonlinear registration and projection onto an alignment-invariant tract representation (Andersson, Jenkinson, Smith, & Andersson, 2007a, 2007b). This technique solves the issue of voxel alignment between participants, ensuring that only voxels that are present in all subjects are included, and does not require smoothing. All FA images were then aligned to the 1x1x1mm FMRIB58_FA standard space target, using a nonlinear registration (Andersson et al., 2007a, 2007b). A standard-space version of each subjects FA image was generated by transforming the FA image to MNI152 standard space (Rueckert et al., 1999). An image containing the mean FA values for all of the subjects was generated and projected onto a mean FA skeleton image, where the skeleton is representative of the centers of all of the tracts common to the subjects. Lastly, for each subject, FA data were projected onto the mean FA skeleton to carry out voxel-wise cross-subject analyses.

### 2.5 Probabilistic tractography

We used the FMRIB linear image registration tool (FLIRT) to register the diffusion-weighted images to the skull-stripped T1 structural image and to a standard MNI152 brain image. Probabilistic tractography was then conducted to estimate the most likely pathway from the seeded regions located in the CR/AF of both the left and right hemispheres (Behrens et al., 2003, 2007). These four seeds were derived by tracing a 2x3 voxel (CR) or 2x2 voxel (AF) seed for each participant in their individual diffusion space for each hemisphere. To ensure that tracing was consistent across subjects, we first selected the most medial slice of their diffusion image in the sagittal view. From this image, we selected the midpoint of the corpus callosum, and then moved laterally into the adjacent CR to locate the seed space. We used individual FA maps for each subject to ensure the six-voxel seed was placed within a section of the tract that diffused in the ventral-dorsal orientation. To locate the seed space for the AF, we centered the FA maps on the middle of the tract (determined by Vandermosten et al. as −32, −24, 22mm), and subsequently adjusted the location to ensure the seed was drawn in a region where the tracts diffused in the anterior-posterior orientation. Finally, we seeded the tractography from these regions 5000 times to calculate the most probable tracts, and only included voxels for which at least 1200 streamlines passed through the resulting tracts.

The *a priori* selection of seed regions makes this type of analysis hypothesis rather than data driven, with the additional benefit of increased statistical power compared with whole-brain analysis. This is because tractography is conducted on a per-subject basis using non-normalized brain data, to allow taking into account individual differences in anatomy, which are often substantial. In addition, probabilistic tractography allows tracking white matter pathways in a way that overcomes some of the shortcomings associated with traditional deterministic tractography (Behrens et al., 2007; Ciccarelli et al., 2006). For example, one well-known issue with deterministic tractography is that it cannot track into regions that are highly curved or display low FA, and has difficulties resolving crossing fibers. The probabilistic algorithm solves these problems by modeling the uncertainty of fiber direction—in any region where direction of the tract is not clear, it tracks white matter in multiple directions, and weights each direction by its probability estimate. In the present study, we used this method to allow for more flexible and reliable tracking of white matter pathways, through regions known to branch in multiple directions and curve to a high degree.

Statistical analyses were performed in R (R Core Team, 2016) and JASP (JASP Team, 2016). A list of all CRAN packages used for our analyses is available in the online material. In this Results section, we report Bayesian model comparisons, to allow quantifying the degree of evidence for a given model compared to other models tested, as well as Bayesian parameter estimations when relevant. Additional information can be found in the online material at https://github.com/davidmoreau/2018_NeuroImage_Clinical

## 3. Results

### 3.1 TBSS

Separate TBSS analyses were run to determine if there were any regions of white matter that differed in connectivity between the groups, controlling for age. Besides comparing all groups with each other, we also contrasted an aggregate of all individuals with a learning disability (dyslexics, dyscalculics, and comorbids) against those without (controls), an aggregate of all individuals with a math difficulty (dyscalculics and comorbids) against those without (dyslexics and controls); and finally a contrast between those with a reading difficulty (dyslexics and comorbids) and those without (dyscalculics and controls). The results of all contrasts showed no difference in FA between the groups, after correcting for multiple comparisons (FWE-corrected, see Figure 1). Uncorrected FA maps are provided in the Supplemental Material (Figure SM1-SM10). Because using a whole-brain approach might not be sensitive enough to detect white matter microstructural differences, we followed these analyses with focused, ROI-based probabilistic tractography.

**Figure 1.**
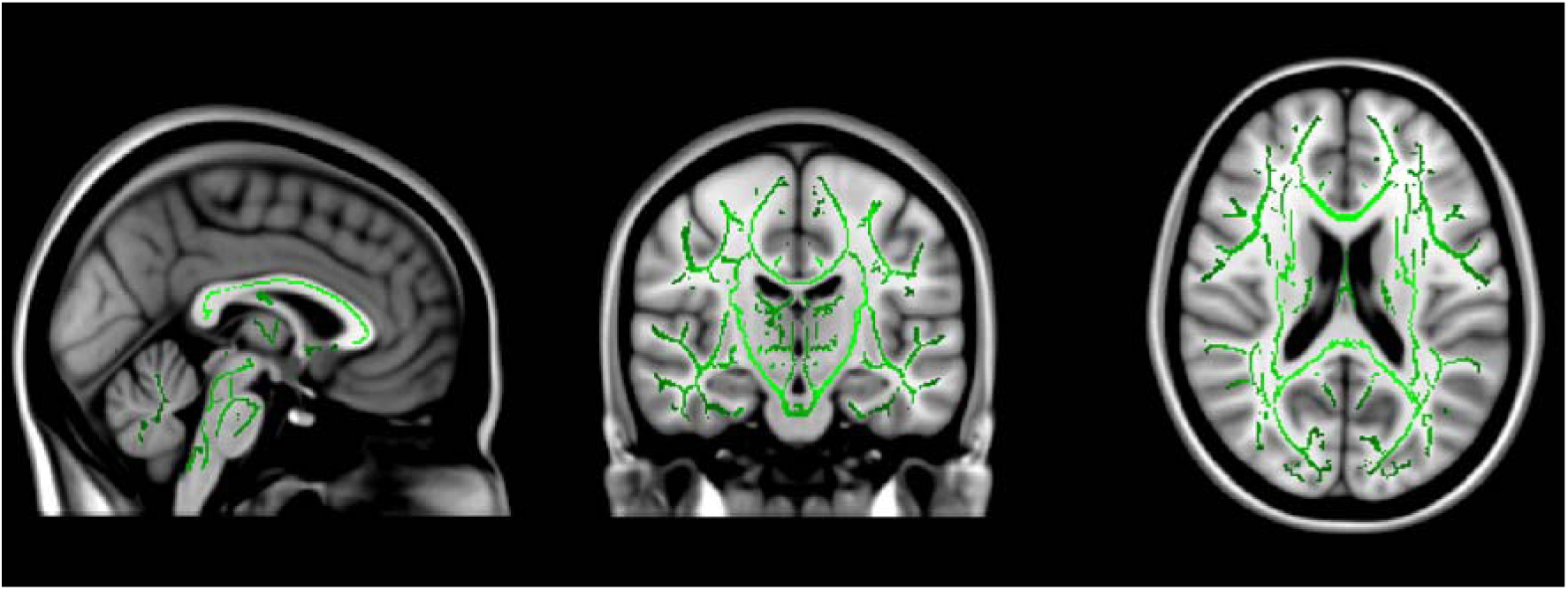
White matter structures showing no regions with significantly higher FA in controls compared to the clinical groups (*p* < 0.05, corrected for multiple comparisons). Green voxels show the mean FA white matter skeleton for these participants. Template image is the standard MNI152 brain.

### 3.2 Probabilistic tractography

We conducted probabilistic tractography on two white matter tracts: the corona radiata and the arcuate fasciculus^1^. We report hereafter Bayesian hypothesis testing and parameter estimation for both tracts, in each hemisphere.

#### 3.2.1 Corona radiata

Two separate Bayesian ANCOVAs with *Mean FA values* as a dependent variable, *Age* and *Gender* as covariates, and *Group* (Control, Dyslexia, Dyscalculia, Comorbid) as a fixed factor showed moderate evidence for the null model (i.e., no difference between groups) over the alternative for both hemispheres [Left: BF_M_ = 4.04, *p*(M | Data) =.37; Right: BF_M_ = 4.42, *p*(M | Data) =.39; Figure 2]. The prior scale was 0.5 for fixed effects, 1 for random effects, and 0.354 for covariates (Morey & Rouder, 2015).

**Figure 2.**
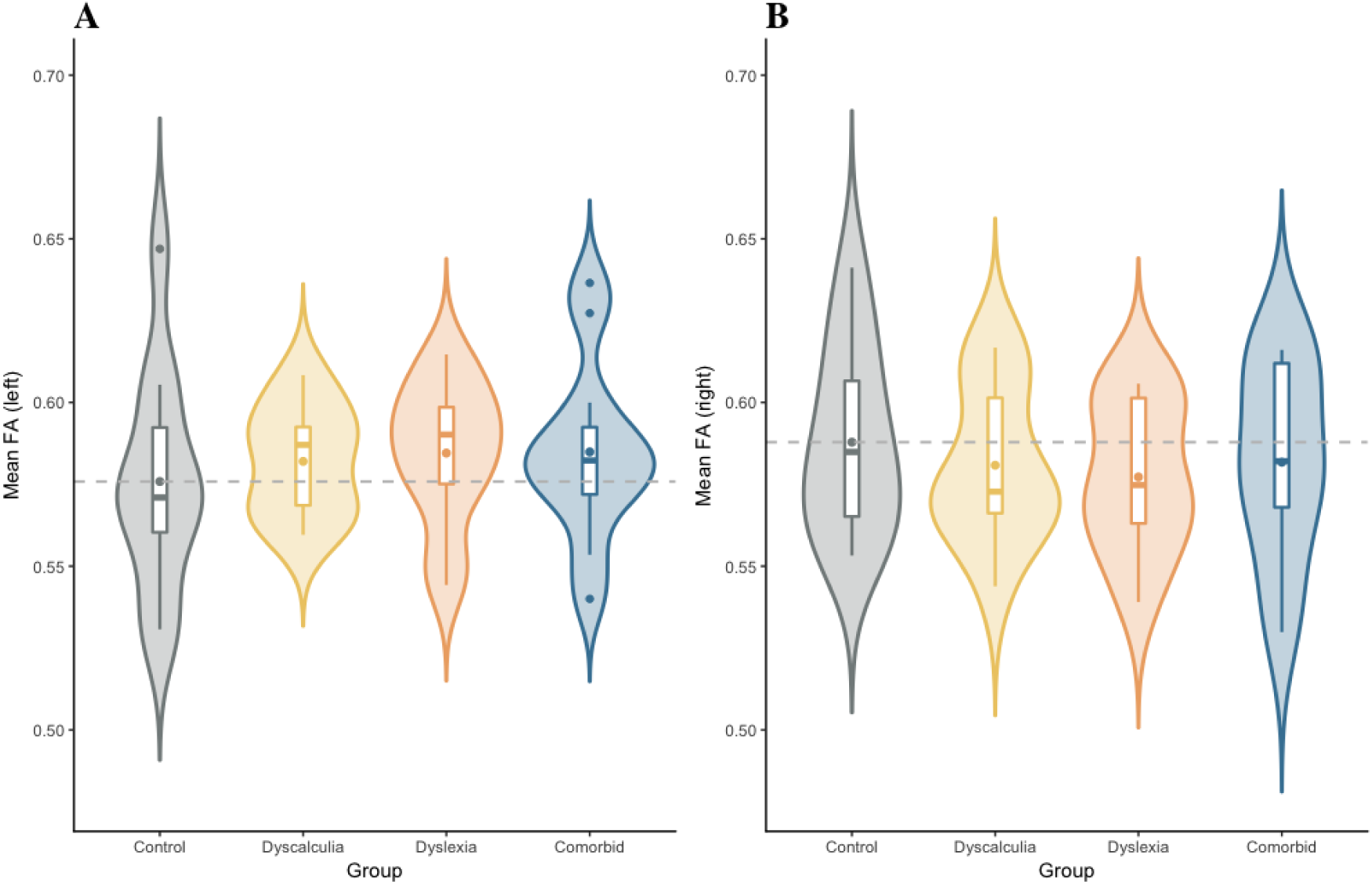
Descriptive plots of FA values across the corona radiata for all four groups (Control, Dyscalculia, Dyslexia, Comorbid). For each group, the plot shows the distribution of gain scores (violin) together with the mean (box central dot), median (box central line), first and third quartile (box edges), minimum and maximum (whiskers), and outliers (outside dots). The dashed line references the mean value for the control group. (A) Left hemisphere. (B) Right hemisphere.

Because differences in the CR have been more frequently associated with reading than mathematical difficulties, we further compared all dyslexics (i.e., dyslexia and comorbid groups) with all non-dyslexics (i.e., dyscalculia and control groups). A Bayesian *t*-test with a Cauchy prior width of 0.95 (i.e., half of parameter values lies within the interquartile range – 0.95 – 0.95; Rouder, Speckman, Sun, Morey, & Iverson, 2009) showed moderate evidence for the null model of no differences between groups (Left: BF_01_ = 3.34, *ε* = 0.029%, Right: BF_01_ = 3.67, *ε* = 0.032%; see Figure 3A-B for prior and posterior distributions, see Figure 4 for the trace plot and marginal density of the posterior *β* distribution). Bayes Factor robustness analyses showed stronger evidence for the null hypothesis with wider Cauchy priors, indicating that our conclusions do not hinge upon a restricted range of priors (Figure 3C-D). Note that the interpretation of the results does not change with normal or *t* distributions to model the prior (see Supplemental Material). Finally, a sequential analysis showed that our data ubiquitously favored the null hypothesis with increasing observations (Figure 3E-F).

**Figure 3.**
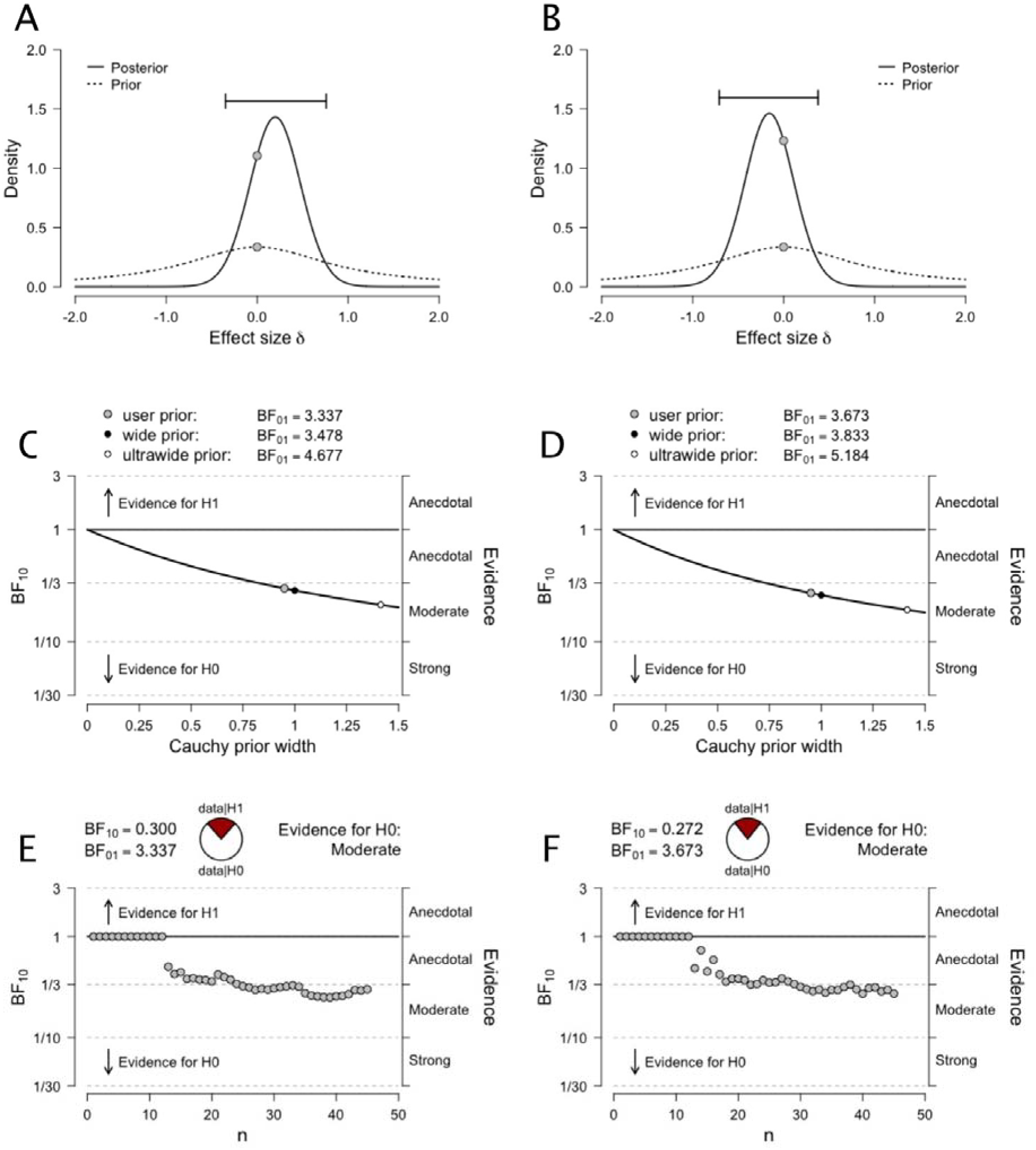
(A-B) Prior and posterior distributions for the comparison between *Groups* (Dyslexics vs. Non-dyslexics) for the left and right CR mean FA values. (C-D) Bayes factor robustness analyses for the comparison between *Groups* (Dyslexics vs. Non-dyslexics) for the left and right CR mean FA values. The figure shows our default prior (gray dot), as well as wide and ultrawide priors (black and white dots, respectively). (E-F) Sequential analysis showing the strength of evidence (as expressed by BF_10_) as *N* increases, for the left and right CR mean FA values. Note that data points are ordered by group.

**Figure 4.**
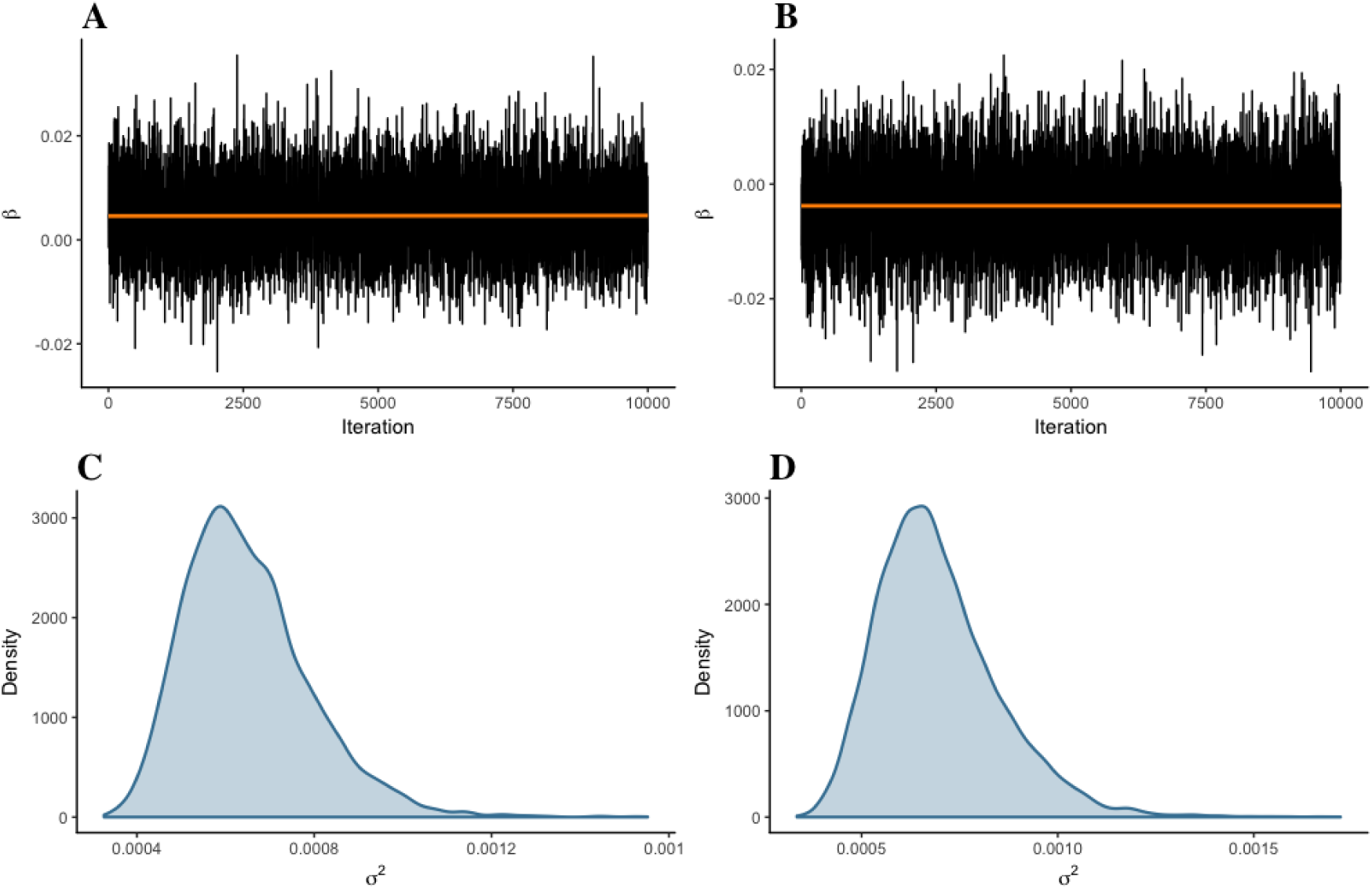
Trace plot and marginal density for parameter *β* of the difference between groups (Dyslexics vs. Non-dyslexics) in mean FA values. (A-C) Left CR. (B-D) Right CR. Estimates were generated from 10,000 iterations, in one chain, with thinning interval of one (i.e., no data point discarded). Both trace plots (the values parameter *β* took during the runtime of the chain) and marginal densities (smoothed histograms of parameter *β* values) indicated that the difference between groups was practically null.

**Figure 5.**
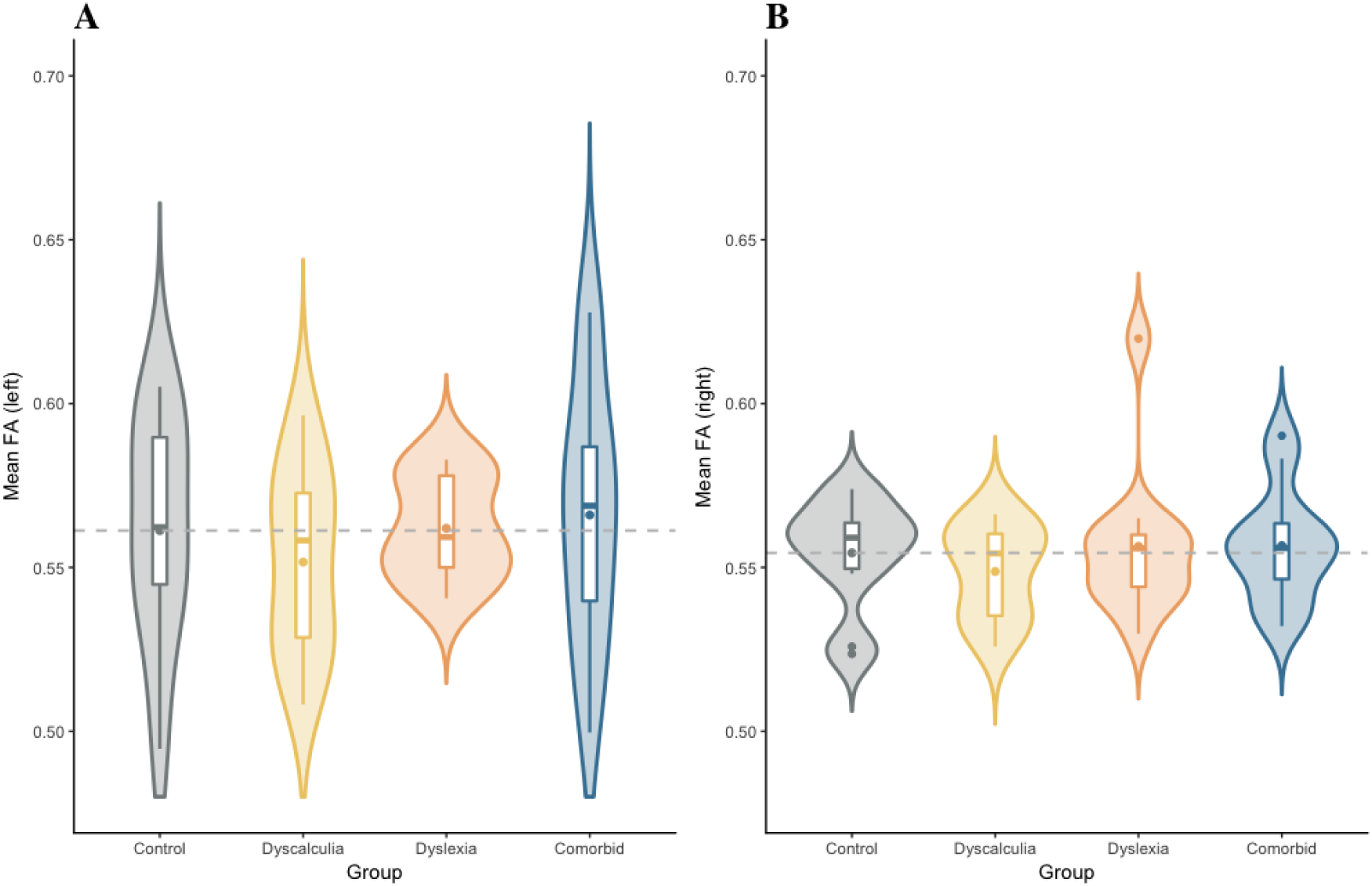
Descriptive plots of FA values across the arcuate fasciculus for all four groups (Control, Dyscalculia, Dyslexia, Comorbid). For each group, the plot shows the distribution of gain scores (violin) together with the mean (box central dot), median (box central line), first and third quartile (box edges), minimum and maximum (whiskers), and outliers (outside dots). The dashed line references the mean value for the control group. (A) Left hemisphere. (B) Right hemisphere.

For transparency, we also contrasted all dyscalculics (i.e., dyscalculia and comorbid groups) vs. all non-dyscalculics (i.e., dyslexia and control groups). Consistent with previous comparisons, a Bayesian *t*-test with a Cauchy prior width of 0.95 (Rouder et al., 2009) showed moderate evidence for the null model of no differences between groups (Left: BF_01_ 3.98, *ε* = 0.033%; Right: BF_01_ = 4.28, *ε* = 0.035%).

#### 3.2.2 Arcuate fasciculus

Two separate Bayesian ANCOVAs with *Mean FA values* as a dependent variable, *Age* and *Gender* as covariates, and *Group* (Control, Dyslexia, Dyscalculia, Comorbid) as a fixed factor showed weak evidence for the null model (i.e., no difference between groups) over the alternative for both hemispheres [Left: BF_M_ = 2.41, *p*(M | Data) =.26; Right: BF_M_ = 2.40, *p*(M | Data) =.26]. The prior scale was 0.5 for fixed effects, 1 for random effects, and 0.354 for covariates (Morey & Rouder, 2015).

Similar to the rationale for testing further differences on the CR tract, and again because differences in the AF have been more frequently associated with reading than mathematical difficulties, we also compared all dyslexics (i.e. dyslexia and comorbid groups) with all non-dyslexics (i.e. dyscalculia and control groups). A Bayesian *t*-test with a Cauchy prior width of 0.95 (Rouder et al., 2009) showed moderate evidence for the null model of no differences between groups (Left: BF_01_ = 3.20, *ε* = 0.029%, Right: BF_01_ = 2.98, *ε* = 0.027%; see Figure 6A-B for prior and posterior distributions, see Figure 7 for the trace plot and marginal density of the posterior *β* distribution). Bayes Factor robustness analyses showed stronger evidence for the null hypothesis with wider Cauchy priors, indicating that our conclusions do not hinge upon a restricted range of priors (Figure 6C-D). Note that the interpretation of the results does not change with normal or *t* distributions to model the prior (see Supplemental Material). Finally, a sequential analysis showed that our data ubiquitously favored the null hypothesis with increasing observations (Figure 6E-F).

**Figure 6.**
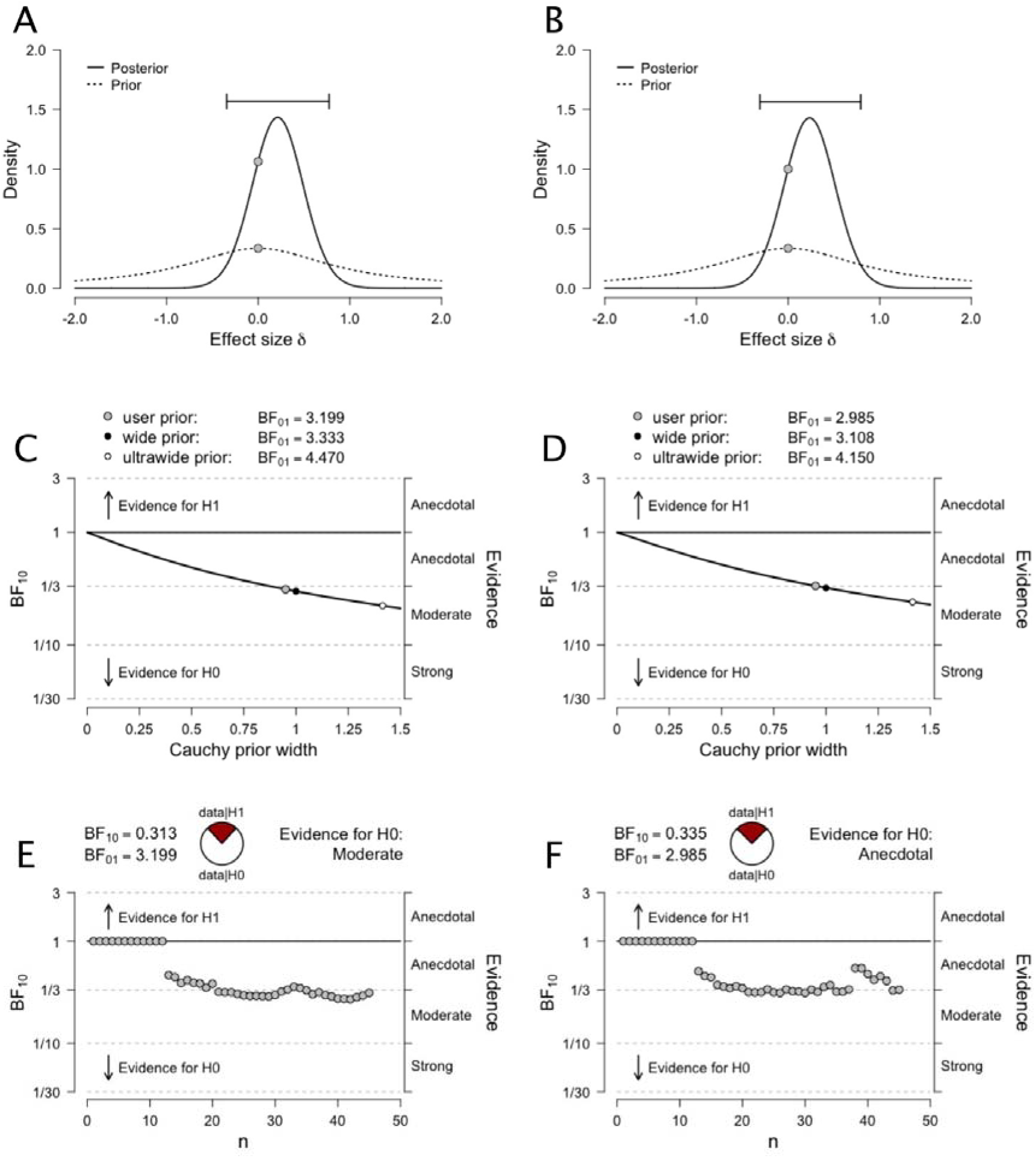
(A-B) Prior and posterior distributions for the comparison between *Groups* (Dyslexics vs. Non-dyslexics) for the left and right AF mean FA values. (C-D) Bayes factor robustness analyses for the comparison between *Groups* (Dyslexics vs. Non-dyslexics) for the left and right AF mean FA values. The figure shows our default prior (gray dot), as well as wide and ultrawide priors (black and white dots, respectively). (E-F) Sequential analysis showing the strength of evidence (as expressed by BF_10_) as *N* increases, for the left and right AF mean FA values. Note that data points are ordered by group.

**Figure 7.**
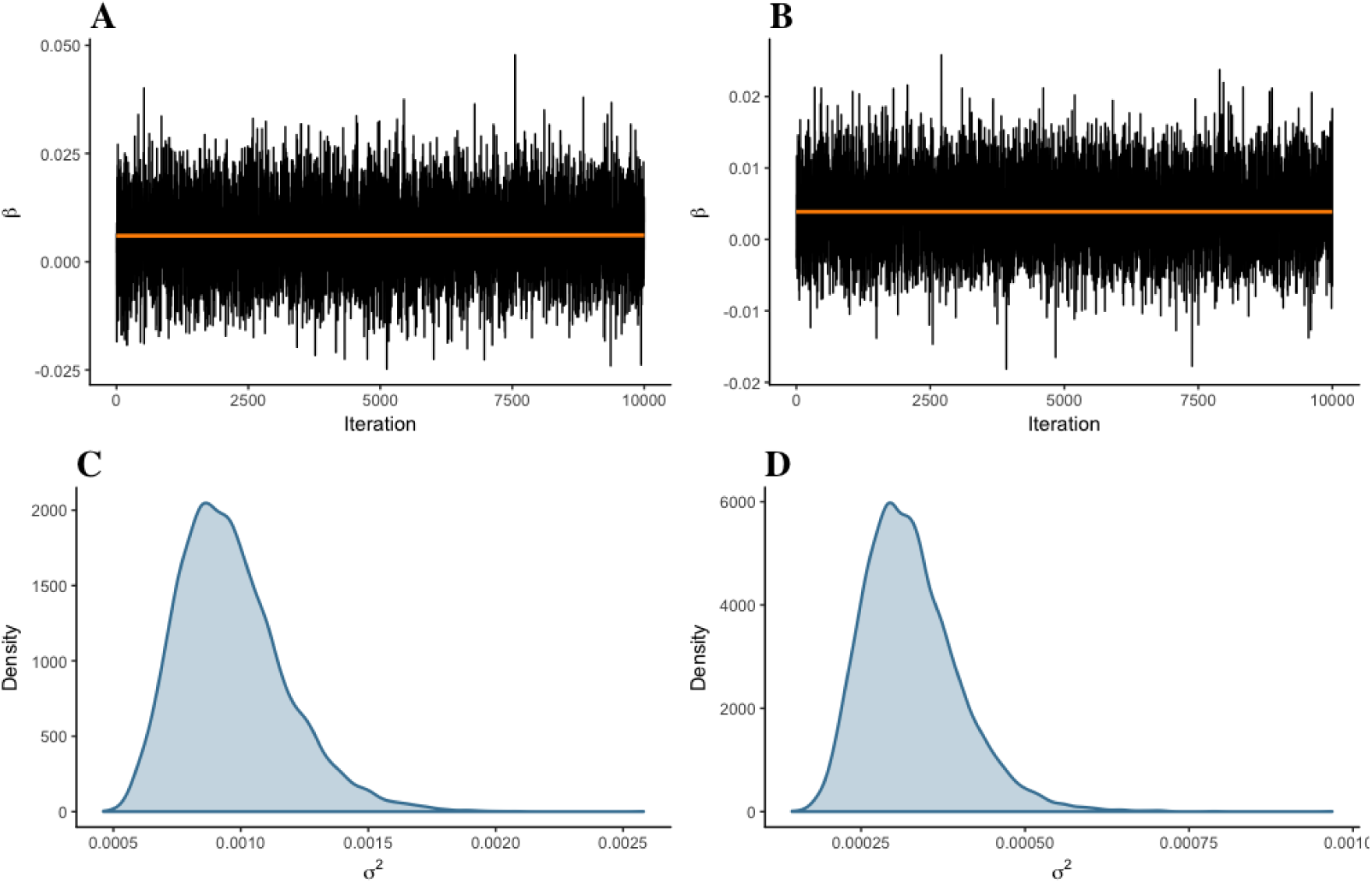
Trace plot and marginal density for parameter *β* of the difference between groups (Dyslexics vs. Non-dyslexics) in mean FA values. (A-C) Left AF. (B-D) Right AF. Estimates were generated from 10,000 iterations, in one chain, with thinning interval of one (i.e., no data point discarded). Both trace plots (the values parameter *β* took during the runtime of the chain) and marginal densities (smoothed histograms of parameter *β* values) indicated that the difference between groups was practically null.

Consistent with our analyses of the CR, we also contrasted all dyscalculics (i.e., dyscalculia and comorbid groups) vs. all non-dyscalculics (i.e., dyslexia and control groups). Consistent with previous comparisons, a Bayesian *t*-test with a Cauchy prior width of 0.95 (Rouder et al., 2009) showed moderate evidence for the null model of no differences between groups (Left: BF_01_ = 4.20, *ε* = 0.035%; Right: BF_01_ = 3.92, *ε* = 0.033%).

### 3.3 Frequentist analyses

We present Bayesian hypothesis and Bayesian parameter estimation for the tractography analyses. Because we understand that the reader may wish to compare these with the frequentist equivalents, we provide these hereafter. Note, however, that because our data provide various degree of support for the null hypothesis of no effect, comparison across frameworks is limited, with inherently different interpretations of null effects.

#### 3.2.1 Corona radiata

Two separate ANOVAs with *Mean FA values* as a dependent variable and *Group* (Control, Dyslexia, Dyscalculia, Comorbid) as a fixed factor showed no significant effect for both hemispheres [Left: *F*(3, 41) = 0.296, *p* =.828, *ω^2^* = 0; Right: *F*(3, 41) = 0.307, *p* =.820, *ω^2^* = 0; Figure 2]. An independent samples *t*-test comparing all dyslexics (i.e., dyslexia and comorbid groups) vs. all non-dyslexics (i.e. dyscalculia and control groups) showed no significant effect for both hemispheres [Left: *t*(42.37) = 0.771, *p* =.446, *d* = 0.23, 95%CI = −0.36; 0.81; Right: *t*(42.95) = −0.613, *p* =.543, *d* = −0.18, 95%CI = −0.77; 0.40]. An independent samples *t*-test comparing all dyscalculics (i.e., dyscalculia and comorbid groups) vs. all non-dyscalculics (i.e., dyslexia and control groups) showed no significant effect for both hemispheres [Left: *t*(40.12) = 0.439, *p* =.446, *d* = 0.13, 95%CI = −0.46; 0.71; Right: *t*(42.96) = −0.160, *p* =.873, *d* = −0.05, 95%CI = −0.63; 0.54].

#### 3.3.2 Arcuate fasciculus

Two separate ANOVAs with *Mean FA values* as a dependent variable and *Group* (Control, Dyslexia, Dyscalculia, Comorbid) as a fixed factor showed no significant effect for both hemispheres [Left: *F*(3, 41) = 0.431, *p* =.732, *ω^2^* = 0; Right: *F*(3, 41) = 0.453, *p* =.717, *ω^2^* = 0; Figure 2]. An independent samples *t*-test comparing all dyslexics (i.e., dyslexia and comorbid groups) vs. all non-dyslexics (i.e. dyscalculia and control groups) showed no significant effect for both hemispheres [Left: *t*(42.66) = 0.831, *p* =.411, *d* = 0.25, 95%CI = −0.34; 0.83; Right: *t*(41.08) = 0.928, *p* =.359, *d* = 0.28, 95%CI = −0.31; 0.86]. An independent samples *t*-test comparing all dyscalculics (i.e., dyscalculia and comorbid groups) vs. all non-dyscalculics (i.e., dyslexia and control groups) showed no significant effect for both hemispheres [Left: *t*(40.82) = −0.271, *p* =.788, *d* = −0.08, 95%CI = −0.66; 0.50, Right: *t*(40.33) = −0.475, *p* =.638, *d* = −0.14, 95%CI = −0.73; 0.44].

## 4. Discussion

The current study aimed to investigate potential differences in white matter between dyslexics, dyscalculics, comorbid individuals, and controls. We used TBSS and probabilistic tractography to test whether there were any reliable differences in FA between the groups of interest. TBSS analyses revealed no meaningful differences in FA between these groups, with none of all the possible single-group contrasts being significant. These analyses were followed by comparisons between (a) an aggregate of all individuals with a learning disability (dyslexics, dyscalculics, and comorbids) and those without (controls); (b) an aggregate of all individuals with a math difficulty (dyscalculics and comorbids) and those without (dyslexics and controls); and (c) between those with a reading difficulty (dyslexics and comorbids) and those without (dyscalculics and controls). Each of these comparisons failed to reveal meaningful differences in FA between any of the groups.

Because it is carried out on group-normalized data, TBSS does not have the same precision as tractography to explore theoretically motivated white matter differences between groups. Therefore, we followed the TBSS analyses with hypothesis-driven tractography analyses, centered on the CR and the AF. In previous work, FA in these tracts has been linked to reading performance and to mathematics performance, making these plausible candidates for reliable white matter differences across our experimental groups. Consistent with the TBSS analysis, however, we found no reliable difference in FA between these groups within the CR and the AF of either hemisphere. Importantly, we used Bayesian hypothesis testing to quantify evidence for the null hypothesis of no difference; this framework allowed us to express a degree of confidence, quantified probabilistically, for the null model of no group difference. This is critical in the present study because evidence for the null hypothesis cannot be attributed to low statistical power, as it also affects evidence for the null; this property was illustrated by sequential analyses for Bayesian inference on each tract. Taken together, the results of both types of analyses therefore support the hypothesis of no systematic difference in FA between the groups of interest.

Our findings contrast with previous research in dyslexia that has reported white matter integrity differences between dyslexics and typical readers. Here, the absence of a difference in FA between individuals with and without dyslexia (comorbids and dyslexics vs. dyscalculics and controls) may cautiously be interpreted as support for the hypothesis of no underlying difference in white matter connectivity between these groups. Although departing from some of the published literature, our results are in line with some of the prior research examining white matter tracts in dyslexic and typical readers. For example, a study by Koerte et al. (2016) reported no difference between dyslexics and typical readers using various diffusion measures, after correcting for multiple comparisons. In addition, a meta-analysis by Vandermosten and colleagues (2012) yielded two clusters with a significant difference in FA between poor and typical readers. These two clusters, one located in the left temporo-parietal region and the other in the inferior frontal gyrus, were then used as seeds for tractography. The analysis showed that they corresponded to the AF and the CR, and highlighted high discrepancies in previous work, with some studies reporting null results similar to ours regarding the CR. Indeed, both Vandermosten et al. (2012) and Yeatman et al. (2011) failed to demonstrate white matter differences between dyslexic and typical readers in this particular tract.

Moreover, it is possible that some prior results implicating the CR in reading have incorrectly ascribed the significant coordinates to this tract. Keller and Just (2009) found a difference in FA in a cluster thought to be a part of the CR; however, upon conducting fiber tracking it was revealed that the cluster was not in fact a part of the CR, but belonged to a tract that ran horizontally from the superior frontal gyrus to the left paracentral lobule. This finding highlights the difficulty of associating coordinates with a certain tract, and raises the possibility that other studies identifying the CR—especially those that did not use tractography, such as that of Matejko, Price, Mazzocco, and Ansari, (2013) and van Eimeren, Niogi, Mccandliss, Holloway, and Ansari (2008)—could correspond to alternative tracts.

Although it should be acknowledged that the literature related to the neural bases of mathematics and of dyscalculia is not as comprehensive as that relating to dyslexia, several studies have reported positive correlations between mathematic performance and FA in multiple white matter tracts, including the AF and CR, and, to a lesser extent, the SLF (Klein et al., 2013; van Beek, Ghesquière, Lagae, & De Smedt, 2014; van Eimeren et al., 2008). These correlations imply that better mathematic performance is associated with greater FA within these tracts, leading to the hypothesis that there might be FA differences within these tracts between dyscalculics and controls. However, other studies suggest the opposite, with dyscalculia being associated with decreases in FA, sometimes in the same tracts (Richards et al., 2008). Perhaps directly reflecting these discrepancies, and similarly to comparative analyses contrasting dyslexics and controls, our results do not support the hypothesis of a robust association between FA and mathematical ability. We should point out, however, that research in this area relating to dyscalculia is rather scant, and further studies are needed to make stronger claims.

The absence of a difference in FA in the CR/AF in the context of dyscalculia may have several potential explanations, which complement the aforementioned. Although both tracts have been associated with mathematical difficulties (see for a review van Eimeren et al., 2008), the scarcity of research on dyscalculia and its white matter correlates does not allow strong, evidence-based hypotheses about specific white matter tracts. Some evidence points out to the involvement of the CR in finger counting strategies and linking fingers with numerical representations (Noël, 2005), which suggest that the specific strategies used to carry out mathematical tasks may influence the extent to which the CR and other white matter tracts are utilized. As a result of differential strategies, it is therefore possible that more fine-grained designs and analyses could identify the precise structural components underlying differences in behavior, in ways that were not possible in the present study. Moreover, the absence of difference in FA in the CR/AF may be influenced by the age of our sample; studies that have observed a difference in FA in the CR/AF have consisted of either children or young adults. In this regard, we are in agreement with Matejko & Ansari, (2015) who noted that further research is necessary to determine the effects of different factors, such as age, on white matter integrity and its relation to mathematical difficulties.

In line with this idea, a potential limitation of this study is that our participants consisted of adults exclusively, whereas the studies that have identified differences in FA in the CR/AF have mainly involved children (Keller & Just, 2009; Niogi & Mccandliss, 2006, although see Richards et al., 2008 for an example in adult populations). Given that research implementing training strategies has demonstrated that white matter integrity is strengthened with repetitive use (Mackey, Whitaker, & Bunge, 2012; Scholz, Klein, Behrens, & Johansen-Berg, 2009), a line of research that led Matejko and Ansari (2015) to suggest that white matter is likely modified by both education and experience, it is possible that with repetitive use across development and into adulthood the differences observed in the CR/AF in children may not persist. In our view, however, this is also a strength of the present study—our participants were adults whose initial difficulties did not go away with aging, suggesting that they represent a reliable sample of individuals with genuine learning disorders, rather than a sample of individuals with apparent, but possibly transient, difficulties.

Finally, an additional limitation of our research is related to the voxel size and the number of directions used for data collection (30 directions). Low resolution could have affected our ability to detect differences, and therefore could be a factor in finding null effects. Furthermore, the ability to resolve the issue of curvature that arises in tractography is reduced with fewer directions; this is typically known as the problem of crossing fibers versus kissing fibers. This is in addition to DTI being a proxy measure of white matter integrity, and to FA being an indirect way to assess connectivity.

Together, these findings nuance previous research in this area, which has reported differences in FA between these groups. Although this could be interpreted as undermining the assumption that anatomical differences underpin the functional and behavioral differences evident between these groups, this would be an exaggeration—it is possible that more subtle structural differences, such as white matter microstructures or gray matter volume, would show different results. More generally, and with the necessary caution required, the present study adds to the growing body of knowledge relating to the etiology of dyslexia, dyscalculia and the comorbidity of these two neurodevelopmental disorders.

## Acknowledgements

The Auckland Comorbidity Study was funded by a University of Auckland Faculty of Science Development Research Fund grant to KEW and postdoctoral fellow AJW. DM and KEW are supported by philanthropic donations from the Campus Link Foundation, the Kelliher Trust and Perpetual Guardian (as trustee of the Lady Alport Barker Trust). DM is also supported by the Neurological Foundation of New Zealand. We would like to thank all of our participants for donating their time, and the various organizations who helped us with recruitment. Thanks to Professor Ian Kirk for invaluable DTI advice.

## Footnotes

1 At the request of a reviewer, we tested for group differences in two additional tracts: the inferior longitudinal fasciculus (ILF) and the inferior fronto-occipital fasciculus (IFOF). Consistent with the results reported in this section, our analyses showed support for the null hypothesis of no difference between groups (see Supplemental Material for details).

